# High-dose oral pyrophosphate inhibits connective tissue calcification in Abcc6 null mice but affects bone structure

**DOI:** 10.1101/2025.07.16.665176

**Authors:** Ibtesam Rajpar, Nicholas Yancy, Jacob Beiriger, Christina Shao, Syed Naqvi, Deana Mancuso, Fatemeh Niaziorimi, Ryan E. Tomlinson, Koen van de Wetering

**Author notes:** Corresponding Author: Koen van de Wetering 1025 Walnut Street College building, Suite 515 Philadelphia, PA 19107. **Data availability**: raw data is available upon reasonable request. **Declarations**: All authors state that they have no conflicts of interest. **Ethics approval Statement**: Animal studies were approved by the Institutional Animal Care and Use Committee of Thomas Jefferson University in accordance with the National Institutes of Health Guide for Care and Use of Laboratory Animals under approval number 22-06-514.

## Abstract

**Pseudoxanthoma elasticum** is a rare inherited disorder marked by abnormal calcium phosphate deposition in soft connective tissues, particularly the skin, arteries, and eyes. It is caused by inactivating mutations in the *ABCC6* gene, which encodes a hepatic efflux transporter. Loss of ABCC6 function leads to reduced plasma levels of pyrophosphate, a key inhibitor of calcification, thereby promoting ectopic mineralization. Oral pyrophosphate therapy has emerged as a potential treatment, but its effectiveness is uncertain. Most ingested pyrophosphate is hydrolyzed in the gut to inorganic phosphate, which may worsen calcification. Moreover, its impact on mineralized tissues remains largely unexplored. *Abcc6-/-* mice closely mimic human pseudoxanthoma elasticum and are widely used in preclinical studies. Although patients are most concerned about ocular complications, eye calcification is rarely assessed in translational studies using Abcc6-/- mice. Using microcomputed tomography we found that ectopic calcification at the ciliary margin is a reliable marker of ocular disease progression in these mice. Administering pyrophosphate in drinking water at concentrations up to 90 mM did not increase calcification in skin or eyes. However, only very high doses effectively prevented ectopic calcification – doses that would equate to an impractical 2.5 g/kg/day of disodium pyrophosphate in humans. These high doses also led to pyrophosphate accumulation in bone and negatively affected bone structure and strength. **In summary**, only supraphysiological doses of orally administered pyrophosphate inhibited ectopic calcification in *Abcc6-/-* mice, but these doses are not feasible for human use and may compromise bone function. These data are especially important considering the currently ongoing clinical trial evaluating the safety and efficacy of oral pyrophosphate administration as a treatment for pseudoxanthoma elasticum.

**LAY SUMMARY:** Pseudoxanthoma elasticum (PXE) is a rare inherited mineralization disorder caused by the absence of functional ABCC6, a liver-expressed protein. This deficiency leads to reduced plasma levels of pyrophosphate, a key inhibitor of mineralization, resulting in abnormal calcium phosphate deposition in the skin, eyes, and blood vessels. Oral pyrophosphate has been proposed as a therapeutic strategy for PXE, and a clinical trial evaluating its efficacy recently began in France. In a PXE mouse model, we show that only very high oral doses of pyrophosphate are effective, but these doses impair bone quality and are not suitable for human use.

## INTRODUCTION

Pseudoxanthoma elasticum (PXE) is a rare, heritable ectopic calcification disorder, with an estimated incidence of around 1 in 25,000^1,2^. This disease is a cause of significant morbidity, and characterized by the deposition of amorphous calcium phosphate in the skin, eyes, and arteries, with grave clinical consequences^3^. To date, no effective cure has been identified for PXE. Following the initial manifestation of symptoms, which may be as early as in the second decade of the life of PXE patients, calcification in the affected organs worsens progressively throughout life, resulting amongst other symptoms in disfiguring skin lesions, intermittent claudication, and vision loss^4^.

PXE is an autosomal recessive disease caused by mutations in the gene encoding the ATP-binding cassette subfamily C member 6 (ABCC6) transporter protein, an integral membrane protein primarily expressed in hepatocytes^5,6^. In the liver ABCC6 mediates the release of large amounts of adenosine triphosphate – among other nucleoside triphosphates – which outside of hepatocytes is converted to adenosine monophosphate and pyrophosphate (PPi) by the ectonucleotide pyrophosphatase/phosphodiesterase 1(ENPP1)^5^. Notably, ABCC6 is one of the primary regulators of the endogenous mineralization inhibitor – PPi – in blood plasma and responsible for more than 60% of the PPi in the circulation of healthy individuals^7^. Although crucial for extracellular PPi homeostasis, the liver itself is not affected by PXE. The ratio of PPi to Pi (inorganic phosphate) in blood plasma is a strong determinant of aberrant calcification, also in PXE patients^8^. Counterintuitively, mineralized tissues like bone and teeth also contain large amounts of PPi^9^. Intriguingly, two recent studies in mice reported a reduced volume of trabecular bone in ageing Abcc6 knockout animals^10,11^, suggestive of a previously unanticipated role of ABCC6 in bone homeostasis.

The Abcc6-/- mouse is an established model for PXE^12^. These animals develop spontaneous calcification in the elastic fibers of arterial blood vessels, the Bruch’s membrane of the eye, and most remarkably in the blood capsules of the vibrissae in the muzzle skin, which is an early biomarker for disease progression^13^. Compared to wild type mice, plasma calcium and phosphate levels remain unchanged in Abcc6-/- mice, while levels of plasma PPi are significantly reduced, similar to human PXE patients^14,15^. Loss of plasma PPi is thought to be responsible for the pathological calcification phenotype in PXE patients and Abcc6-/- mice^15^. Since reduced ABCC6 activity in the liver results in dramatically reduced plasma PPi concentrations in Abcc6-/- mice and PXE patients, augmentation of this metabolite in the circulation is also expected to reduce connective tissue calcification in mice and humans lacking ABCC6.

Few studies have tested this hypothesis in humans and mice^16–18^. When healthy human volunteers consumed high doses of tetrasodium PPi orally, a transient but significant elevation in blood plasma PPi levels was noted^16^. Further, Abcc6-/- mice supplemented with drinking water containing PPi reduced ectopic calcification of the muzzle skin, and when given daily intraperitoneal injections of PPi, inhibited development of acute cardiac calcification^16,17^. These findings allude to the potential of oral PPi as a cost-effective and practical treatment for ectopic calcification in PXE. Further, a number of preclinical therapies are being tested to increase plasma PPi in PXE patients including recombinant ENPP1 therapy, and TNAP inhibition therapy^19^. Notably, at the time of writing this paper, a phase II, double blinded, randomized clinical trial for the treatment of PXE with oral PPi, was already recruiting patients in France^20^.

Significant gaps in our knowledge about the potential of oral PPi treatment to ameliorate the clinical manifestations in PXE remain to be addressed. Among them are the low bioavailability and short half-life of PPi in plasma. Most of ingested PPi is metabolized to Pi in the gut. We hypothesized that the increased levels of uptake of Pi associated with oral PPi administration, may in fact directly contribute to increased ectopic calcification in PXE. In addition, the effect of administration of high doses of PPi on mineralized tissues such as bone has not been explored.

In the current study we evaluated the effect of high doses of orally administered PPi on 1) ectopic calcification of soft connective tissues, and 2), the structural and biomechanical properties of long bones of Abcc6-/- mice. Finally, we evaluated the potency of oral PPi administration to counteract ectopic mineralization in Abcc6-/- animals on a diet high in phosphate and low in magnesium. Such a diet is used by many research groups to accelerate ectopic mineralization, which allows for a reduction in the time animals need to be treated to pick up therapy effects^21,22^. Results from this study are expected to inform and benefit future translational studies in mice and clinical trials for PXE.

## MATERIALS AND METHODS

### Animals

*Abcc6*-/- mice used for this study were on a C57BL/6J background and kept in accordance with the IACUC of Thomas Jefferson University. Both male and female mice were included in every dataset. Age-matched wild-type mice were included as controls. One group of mice were fed a diet of standard rodent (LabDiet 5010) chow from weaning up to six months of age to euthanasia. A second group of animals was fed a special, chemically defined diet (TD.00442) comprising 0.4% calcium, 0.85% phosphate, 0.04% magnesium and 100 IU/g vitamin D, hereafter referred to as ‘acceleration diet’^23^ directly following weaning until mice were ∼two months old. Experimental mice were provided disodium PPi in drinking water at the following concentrations; 5, 10, 22, 45 and 90 mM. Treatment started directly following weaning when animals were 3 weeks old and lasted for duration of life. PPi-containing water was refreshed twice per week.

### Blood and tissue collection

Blood samples were collected from anesthetized mice immediately prior to euthanasia. Blood was collected by cardiac puncture, into 1 ml syringes containing 25 µl of a 8.6% tripotassium-EDTA solution. Samples were dispensed into Eppendorf tubes and centrifuged at 4000 rpm (∼1500 rcf) for 10 minutes at 4°C. The upper layer of blood plasma was collected into fresh tubes and centrifuged at 8000 rpm for 10 minutes to generate platelet-poor plasma, which was stored at -80°C until analysis.

Directly following blood collection, the following tissues were collected to quantify ectopic calcification: eyes, kidneys and muzzle skin. Muzzle skin was stored at -80°C whereas eyes and kidneys were stored in 70% ethanol after fixation in 10% formalin for 2 days. Femora and tibias were collected and stored in hydrated gauze at -80°C.

### Analysis of bone structure and mechanics

Femora were scanned using a Bruker micro computed tomography (µCT) analyzer with a 1mm aluminum filter^24^. Bone scans were collected using scanning parameters of 55 kV and 181 µA at a resolution of 13 microns. Scans were reconstructed using nRecon (Bruker) software. Cortical and trabecular bone parameters were quantified using CTan (Bruker) software. Bone mechanical properties were analyzed with three-point bending assay using a material testing system (TA Instrument Electroforce 3200 Series III). Femora were placed on custom made fixtures with the condyles facing down and a span length of 7.6 mm was recorded with calipers. Bones were then fractured to failure with a monotonic displacement ramp of 0.1 mm/s. A custom GNU Octave script was used to derive material properties.

### Quantification of ectopic calcification in soft tissues

The left muzzle skin was stored at -80°C until µCT analysis on a SkyScan 1275 (Bruker, Billerica, MA) at 10 µm resolution using settings described earlier for bone^24^. Ectopic mineral deposits were quantified using CT Analyser (Bruker, Billerica, MA), using a lower cut-off value of 55 kV. Eyes and kidneys were first fixed in 10% formalin before scanning. Kidneys and eyes were scanned at a resolution of 10 and 7 µm, respectively. Ectopic mineral deposits in kidneys and eyes were quantified using lower cut-off values of 55 and 65 kV, respectively. Representative images of ectopic mineral deposits in various tissues were generated using DataViewer software (Bruker, Billerica, MA) in combination with Fiji for Maximum Intensity Projection (MIP).

### PPi quantification

For analysis of whole bone PPi (nanomoles/mg weight), first, the epiphyseal region of the femur was removed and one half was centrifuged at 17000 rpm for 1 minute to remove bone marrow. Similarly, tibiae were cut from one end to release the marrow by centrifugation. Marrow-free bones were cleaned of peripheral soft tissue and dissolved in 10% formic acid (1 ml per 25 mg bone) overnight in a shaker-incubator set at 60°C. The next day, bone solutions were centrifuged at 17000 rpm for 10 minutes to remove organic composites, and part of the supernatants were transferred to fresh tubes. Bone supernatants were further diluted in ddH_2_O (1:400 dilution) for assay of PPi. A standard enzymatic assay of ATP sulfurylase was used to convert PPi to ATP for quantification, as per the method described in Jansen *et al.* ^25^. 2.5 µl of sample was incubated with 77.5 ul of a master mix comprising of 80uM magnesium, 75mU/ml ATP sulphurylase, 1µM adenosine 5’ phoshosulphate and 50mM HEPES buffer, for 30 minutes at 30°C and 10 minutes at 90°C. ATP was quantified using the ATP detection reagent SL (BioThema, Stockholm, Sweden). Bioluminescence was measured using a Tecan microplate reader. Sample PPi concentrations were extrapolated from a standard curve of known PPi concentrations and subsequently converted to nmoles PPi/ mg bone.

PPi in plasma was quantified essentially as described in Jansen *et al.* 2014^25^, with a few minor modifications: PPi was converted into ATP using ATP sulfurylase in the presence of excess adenosine 5’ phosphosulfate (APS). To 2.5 μl of sample, 80 μl of a mixture containing 75 mU ATP sulfurylase (New England Biolabs, Ipswich, MA), 3 μmol/L APS (Santa Cruz Biotechnology), 80 μmol/L MgCl_2_ and 40 mmol/L HEPES (pH 7.4) was added. The mixture was incubated for 30 minutes at 37°C, after which ATP sulfurylase was inactivated by incubation at 90°C for 10 min. Generated ATP was subsequently quantified in 10 µl using 30 µl of SL reagent (BioThema, Stockholm, Sweden).

### Statistical analysis

For quantification of hydroxyapatite in non-bone tissues and PPi in bone and plasma, bars in graphical data represent the mean, error bars represent standard error of mean, and dots represent individual data points. Asterisks represent significant differences compared to untreated Abcc6-/- mice. For comparisons of group means, one-way analysis of variance was used with post hoc Dunnett’s tests (p ≤ 0.05). For analysis of bone structure and mechanics, standard linear regression analysis was used, and p-values indicate the significance of group trends. Dotted lines represent 95% confidence intervals, and dots are individual data points. Statistical analysis was carried out in Prism 9 (Graphpad).

## RESULTS

### Supraphysiological doses of oral PPi do not stimulate ectopic calcification in Abcc6-/- mice

Tissues most likely to be affected by ectopic calcification in the Abcc6-/- mouse model were scanned and analyzed. As expected, we observed robust calcification in the blood capsules surrounding the whiskers in the muzzle skin of Abcc6-/- mice receiving drinking water without PPi. Control wild type animals did not exhibit any ectopic mineral deposition in the muzzle skin (Fig. 1A). Muzzle skin calcification in Abcc6-/- mice could only be inhibited by providing drinking water with high concentrations (45 and 90 mM) of PPi, with lower concentrations being completely ineffective. These results are somewhat contrasting previous work where substantial reduction in muzzle skin calcification was seen in Abcc6-/- animals receiving water containing as little as 10 mM PPi^16^. High doses of PPi did however not stimulate ectopic calcification in the muzzle skin either (Fig. 1A), disproving our hypothesis. Most PXE patients are particularly worried about progression of calcification in their eyes, as this results in vision loss, with many patients eventually becoming completely blind. It is therefore remarkable that calcification in the eye is almost never followed in translational studies testing new treatments for this condition. In the present study, we therefore also determined calcification in the eyes of the Abcc6-/- mice. These studies revealed mineral deposition in a distinct region of the eye known as the ciliary margin. Notably, eyes of control wild-type mice were completely devoid of calcification (Fig. 1B). To the best of our knowledge this is the first report showing calcification of this specific structure in the Abcc6-/- mouse eye. We will provide a detailed description of this novel phenotype in a separate paper (manuscript in preparation). Interestingly, similar to muzzle skin, only mice supplemented with a high dose of 90 mM PPi in their drinking water did not develop calcification in their eyes, with drinking water containing 45 mM PPi being only partially effective (Fig. 1B).

**Fig 1.**
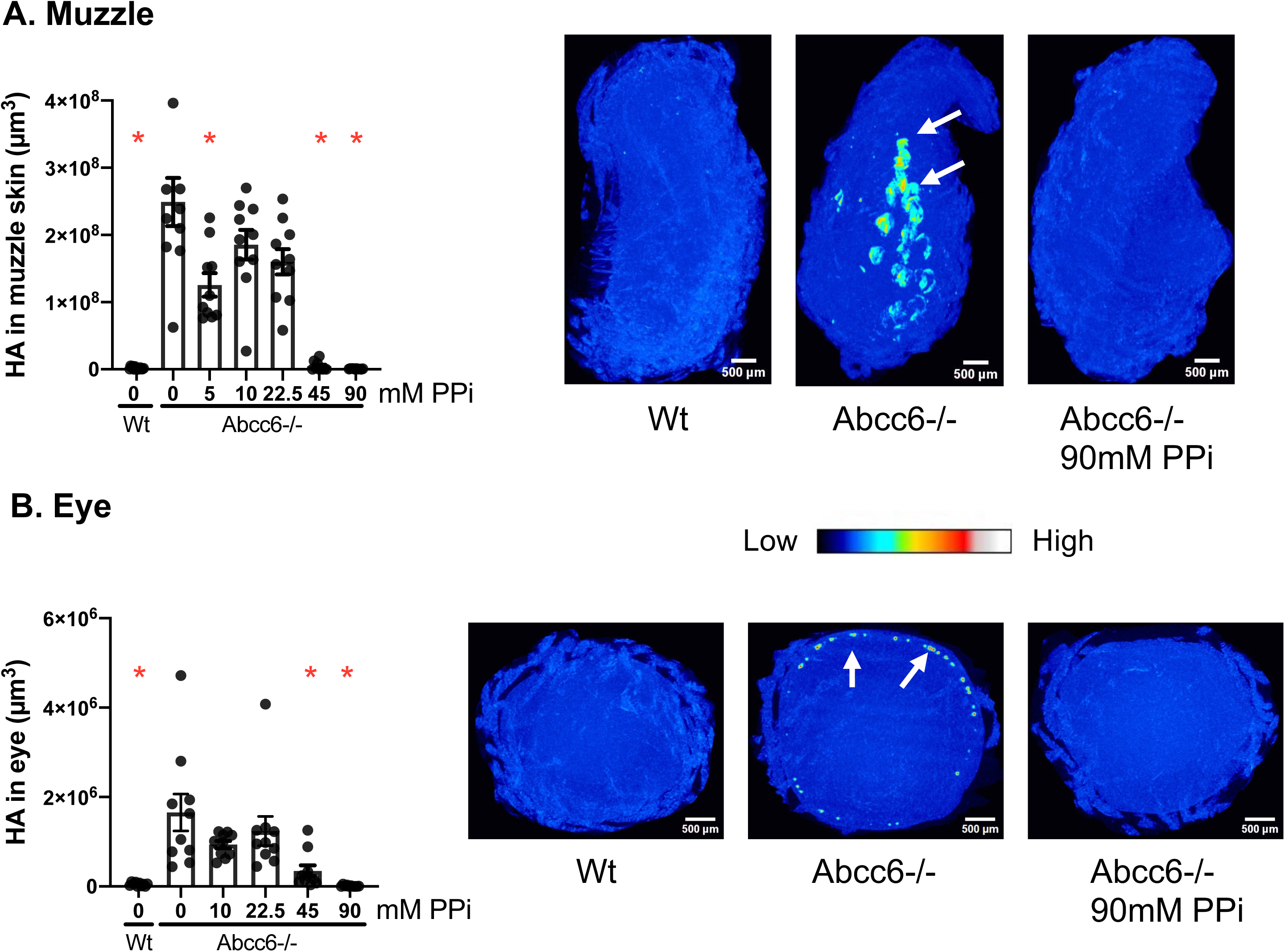
Only high doses of orally administered PPi prevent ectopic calcification in the muzzle skin and eyes of Abcc6-/- mice. Gross examination of microCT sections show presence of mineral deposits (white arrows) in the muzzle skin A) and eye B) of Abcc6-/- animals, but not Wt and Abcc6-/- mice treated with 90mM PPi. Ectopic calcification was quantified using microCT scans of muzzle skin A) and eye B). Group means were compared using one way ANOVA with post hoc Dunnett’s tests. Red asterisks indicate significant differences compared to untreated Abcc6-/- mice, and significance was set at p < 0.05, n=10 per group. Error bars represent SEM.

A specially formulated rodent diet low in magnesium and high in phosphate is used by several research groups to accelerate disease progression in models of ectopic calcification^23^. Such a diet allows for a reduction in the time needed to treat animals in translational studies assessing the efficacy of new experimental therapies. It is unknown, however, if such a diet at the same time affects treatment effectiveness. Possibly, by creating an environment favoring calcification, more potent anti-mineralization strategies are needed. As expected, Abcc6-/- mice receiving the acceleration diet developed robust ectopic calcification in the muzzle skin, already at ∼2.5 months of age (Fig. 2 A,B). Remarkably, under these conditions only drinking water containing 90 mM PPi effectively inhibited muzzle skin calcification in Abcc6-/- mice (Fig. 2A). A specific feature of Abcc6-/- mice receiving the acceleration diet is the development of extensive kidney calcification (Fig. 2B), something not seen in animals fed the standard diet. PPi was unable to prevent the kidney calcification at any of the concentrations tested. Intriguingly, despite creating a pro-calcification state, no ectopic calcification of the ciliary margin of the eye was observed in any of the Abcc6-/- animals maintained on acceleration diet. In summary, PPi was less effective in preventing ectopic calcification in Abcc6-/- mice receiving the acceleration diet.

**Fig 2.**
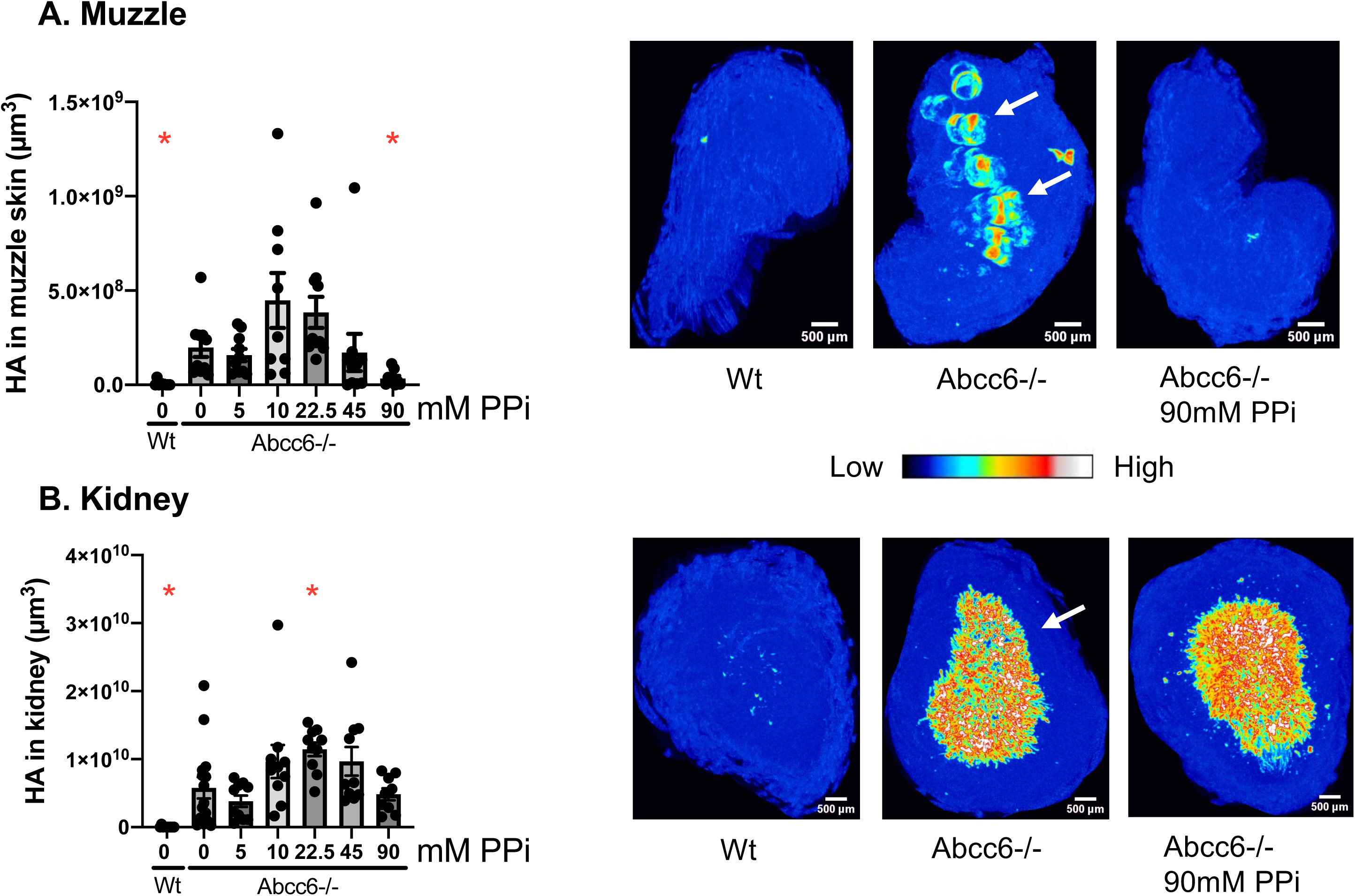
PPi administered via the drinking water reduces ectopic calcification of muzzle skin but not kindeys of Abcc6-/- mice fed a diet accelerating calcification. Gross examination of microCT sections show presence of mineral deposits (white arrows) in the muzzle skin A) and kidneys B) of untreated Abcc6-/- animals, and in the kidneys B) of 90 mM PPi-treated Abcc6-/- animals. Group means were compared using one way ANOVA with post hoc Dunnett’s tests. Red asterisks indicate significant differences compared to untreated Abcc6-/- mice, and significance was set at p < 0.05, n=10 per group. Error bars represent SEM.

Variability in the volume of ectopic mineral deposition in organs and tissues of untreated Abcc6-/- mice analyzed in this study, namely muzzle skin, eyes and kidneys, was relatively large, something previously reported by other laboratories^26,27^. This is a remarkable observation given that knockout mice were bred on the same homogeneous genetic background (C57Bl/6J), fed a similar diet and often were littermates. We will come back to this observation in the discussion section.

### Orally administered PPi accumulates in long bones of Abcc6-/- mice

ABCC6 is responsible for 60-70% of the PPi found in the systemic circulation^15,25^. Reduced plasma PPi is one of the demarcating features of Abcc6-/- mice^15^. Results from this study agree with this hypothesis (Supp Fig. 1A, B). Specifically, Abcc6-/- mice had plasma PPi levels that were 60-70% lower than the levels found in age-matched WT mice (Supp Fig. 1A, B). Even at the highest dose tested of 90 mM, we did not detect increased levels of plasma PPi in Abcc6-/- animals receiving PPi via their drinking water. Lack of an increase in plasma PPi as a result of oral PPi supplementation has been reported before^16^ and likely reflects the transient nature of increases in plasma PPi concentrations following consumption of PPi-treated drinking water. Interestingly, diet did affect plasma PPi concentrations with WT mice fed the standard diet exhibiting much higher plasma PPi levels than WT mice fed the acceleration diet. Likewise, also Abcc6-/- mice had higher plasma PPi levels on a standard diet than when on the acceleration diet (Supp Fig. 1A, B).

PPi tightly binds to mature calcium hydroxyapatite crystals in the bone matrix. Bone therefore contains high levels of PPi, which can be attributed to ANK-mediated ATP efflux from osteoblasts and subsequent hydrolysis by ENPP1 in the extracellular matrix^28,29^. Although accumulation of a mineralization inhibitor in bone seems counterintuitive, PPi has been hypothesized to stabilize existing hydroxyapatite crystals^30^. It is at present unclear how much diffusion from the skeletal vasculature contributes to PPi incorporation into the bone matrix. We therefore examined if supplementation via drinking water increased PPi content of long bones. First, PPi content in femora of WT mice was significantly higher than than in femora of untreated Abcc6-/- mice on both, standard and acceleration diet (Fig. 3A,C), suggesting that some PPi in circulation accumulates in the bone matrix. Intriguingly, Abcc6-/- mice provided with 45 and 90 mM PPi in their drinking water had increased levels of PPi in their femur bone matrix (Fig. 3A). These results from six-month-old mice were confirmed in adolescent mice of ∼10 weeks maintained on acceleration diet (Fig. 3C). Treatment of Abcc6-/- mice with 45 mM PPi resulted in an approximately 30% increase in total matrix PPi (Fig. 3C).

**Fig 3.**
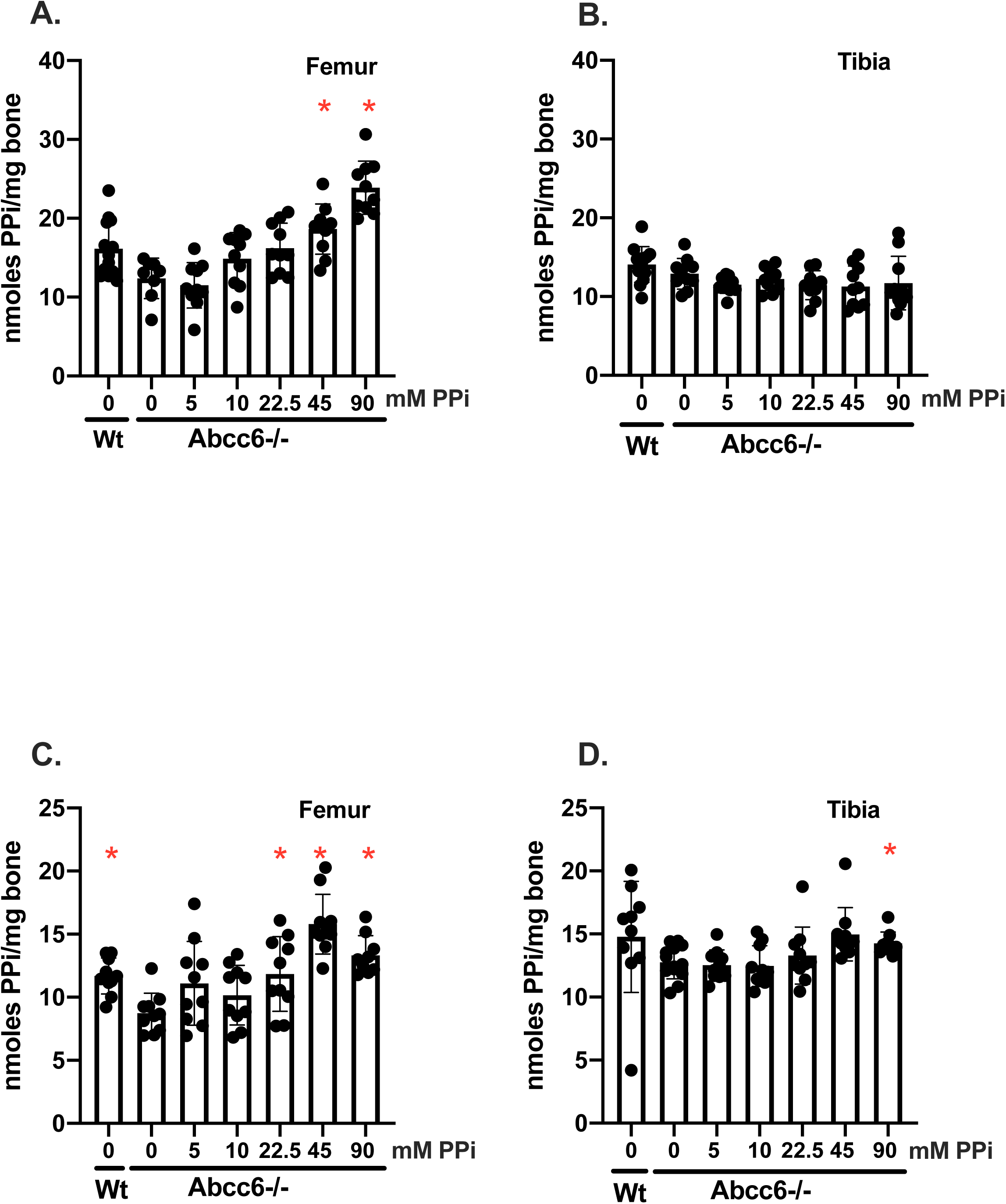
PPi adminstered via the drinking water to Abcc6-/- mice accumulates in long bones. PPi was quantified in femora and tibiae of mice given standard (A, B) and acceleration diet (C, D), and quantified using a standard luciferase assay for ATP. Group means were compared using one way ANOVA with post hoc Dunnett’s tests. Red asterisks indicate significant differences compared to untreated Abcc6-/- mice, and significance was set at p < 0.05, n=10 per group. Error bars represent SEM.

Further, this trend was also seen in tibiae of Abcc6-/- mice maintained on acceleration diet, with increased levels of PPi in bone mineral seen in animals receiving 90mM PPi via their drinking water (Fig. 3D). PPi in the drinking water did not seem to affect accumulation of PPi in tibiae of Abcc6-/- mice maintained on standard diet, however (Fig. 3B).

### Supplementation of Abcc6-/- mice with PPi impacts bone mechanical properties

Our data indicate that oral PPi supplementation does not increase ectopic calcification of soft connective tissues of Abcc6-/- mice. However, much higher doses of PPi were needed to inhibit calcification than previously reported^16^. As high doses of orally administered PPi resulted in the accumulation of this mineralization regulator in bone tissue, we next determined if this negatively affected structure and strength of the long bones. Femora of Abcc6-/- mice were analyzed by microCT, then subjected to three-point bending to quantify bone mechanical properties. In animals fed our standard rodent diet, ultimate moment was reduced in Abcc6-/- mouse femora with increasing oral PPi dose, and this trend was close to being statistically significant (Fig. 4). This decrease suggests that increased PPi incorporation compromises the load-bearing capacity of these femora (Fig. 4). However, other parameters of biomechanical strength and stiffness including bending rigidity, toughness and Young’s modulus were not affected by oral PPi treatment (Fig. 4). PPi treatment also did not result in major differences in the bone mineral density of cortical and/or trabecular bone of the femur (Fig. 5, Supp Fig 3). Small differences in the cross-sectional thickness of the femur (Fig. 5) and thickness of trabeculae (Supp Fig. 3) are noteworthy, but do not appear to strongly affect bone structure. Thus, lower ultimate moment may be caused by changes in the material properties of hydroxyapatite induced by the increased levels of PPi in the femora.

**Fig 4.**
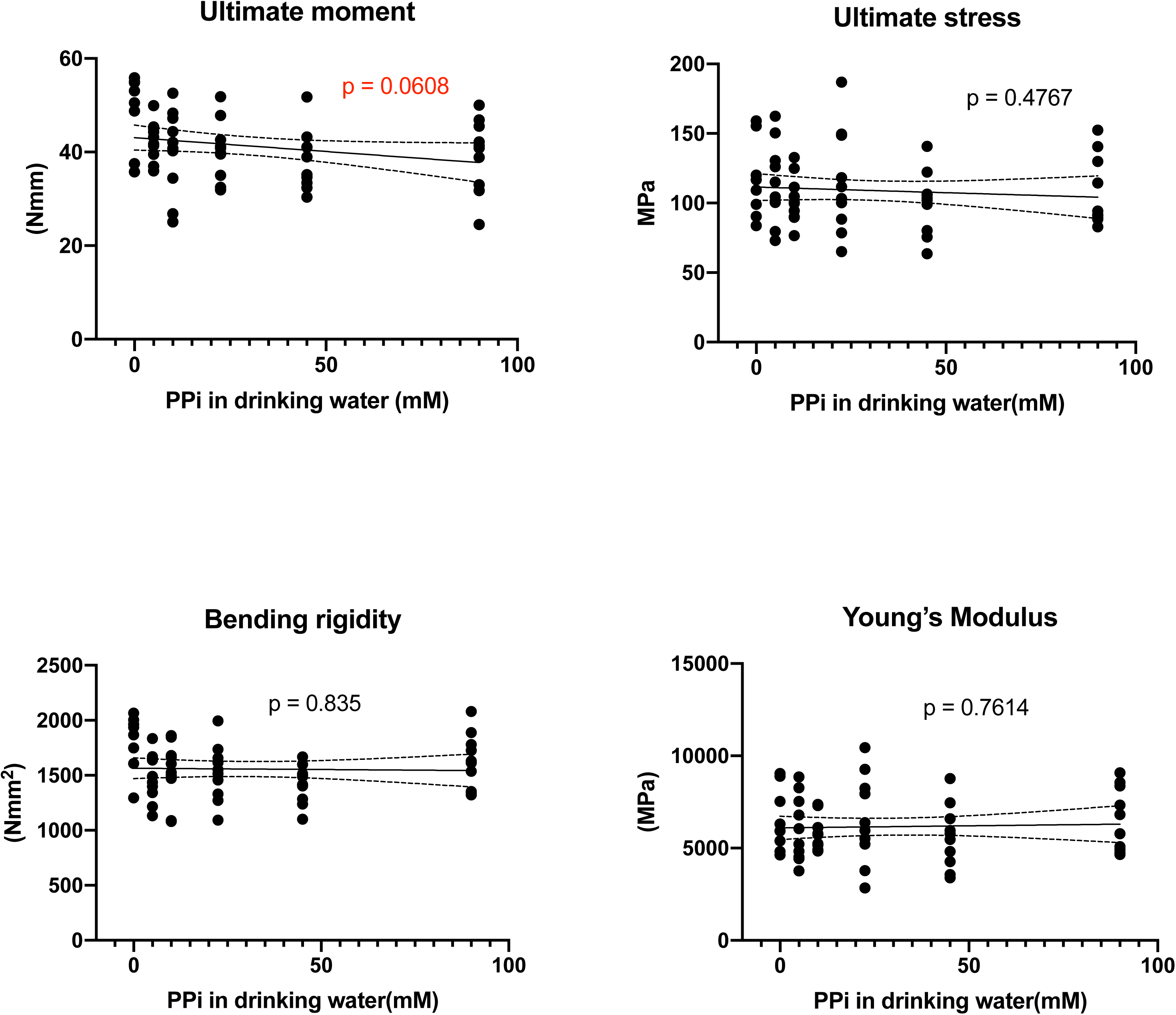
PPi provided via the drinking water affects the mechanical properties of femora from Abcc6-/- mice. Various parameters of bone strength and stiffness were quantified using microCT and the three point bending assay. Group trends were quantified with standard regression analysis. Lines represent 95% confidence intervals. Significance was set at p < 0.05, n=10 per group. Dots are individual data points.

**Fig 5.**
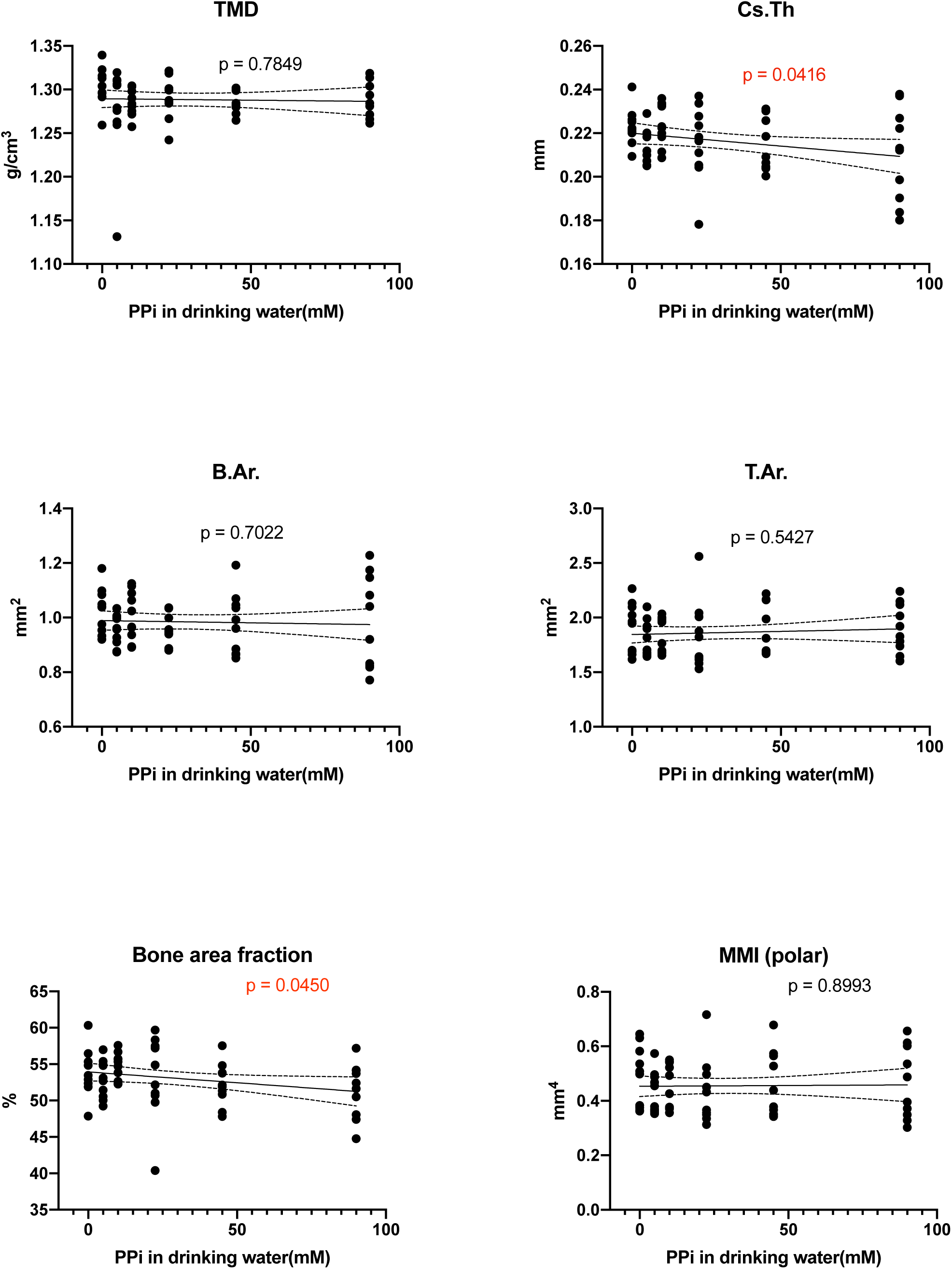
Effect of PPi treatment on the cortical bone properties of femora from Abcc6-/- mice. Various structural parameters of cortical bone were quantified with microCT. TMD (tissue mineral density), Cs.Th (cross sectional thickness), B.Ar. (Bone area), T.Ar. (Tissue area), Bone area fraction (B.Ar./T.Ar.*100), MMI (mean polar moment of inertia). Group trends were quantified with standard regression analysis. Lines represent 95% confidence intervals. Significance was set at p < 0.05, n=10 per group. Dots are individual data points.

In contrast, orally administered PPi had a more dramatic effect on bone structural and mechanical properties in Abcc6-/- mice receiving the acceleration diet (Fig. 6). Significant reductions in bone stiffness and strength were noted when Abcc6-/- mice were treated with drinking water containing PPi (Fig. 6). Specifically, the ultimate moment and bending rigidity of femora was greatly reduced with increasing consumption of PPi via the drinking water (∼2.5 months, p < 0.0001) (Fig. 6). Femora in this group also exhibited significant differences in cortical and trabecular bone structure as observed by microCT analysis. Several markers of cortical bone structure and mechanics, including cortical bone area, mean polar moment of inertia (MMI) and cross-sectional thickness of femora were reduced with increased intake of PPi (Fig. 7). Further, a small but remarkable reduction in trabecular bone mineral density was also noted (p = 0.0548) and appeared to be most pronounced in the 90mM PPi group (Supp Fig. 5). We also noted a loss of trabecular thickness in this group of femora (Supp Fig. 5), but this was not accompanied by a parallel increase in trabecular spacing. Together, these alterations in bone structure may contribute to the reduced femoral strength in Abcc6-/- mice receiving drinking water containing PPi.

**Fig 6.**
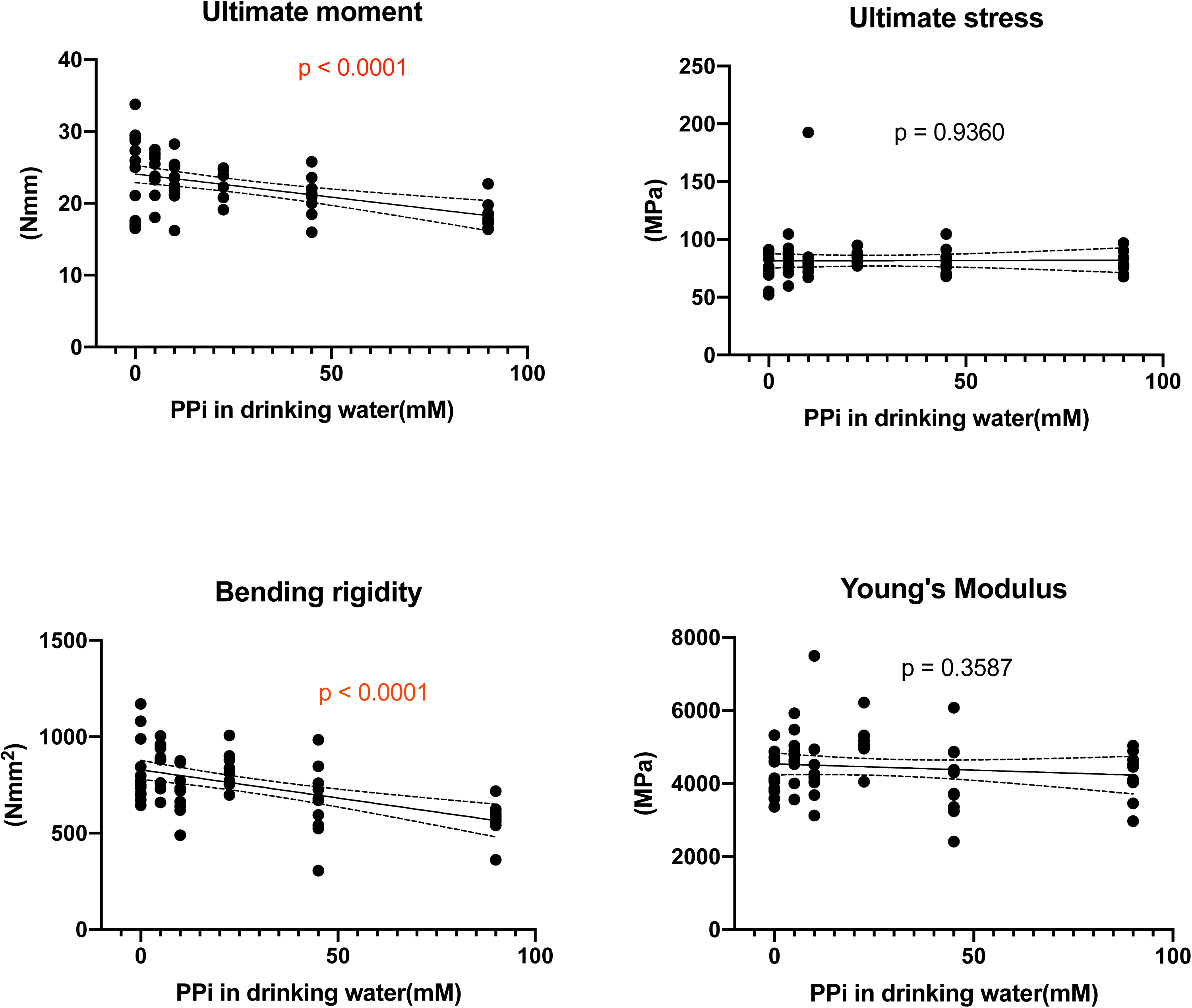
Mechanical properties of femora from Abcc6-/- mice receiving a diet that accelerates ectopic calcification were substantially affected by oral PPi treatment. Various parameters of bone mechanical strength were quantified using microCT and the three point bending assay. Group trends were quantified with standard regression analysis. Lines represent 95% confidence intervals. Significance was set at p < 0.05, n=10 per group. Dots are individual data points.

**Fig 7.**
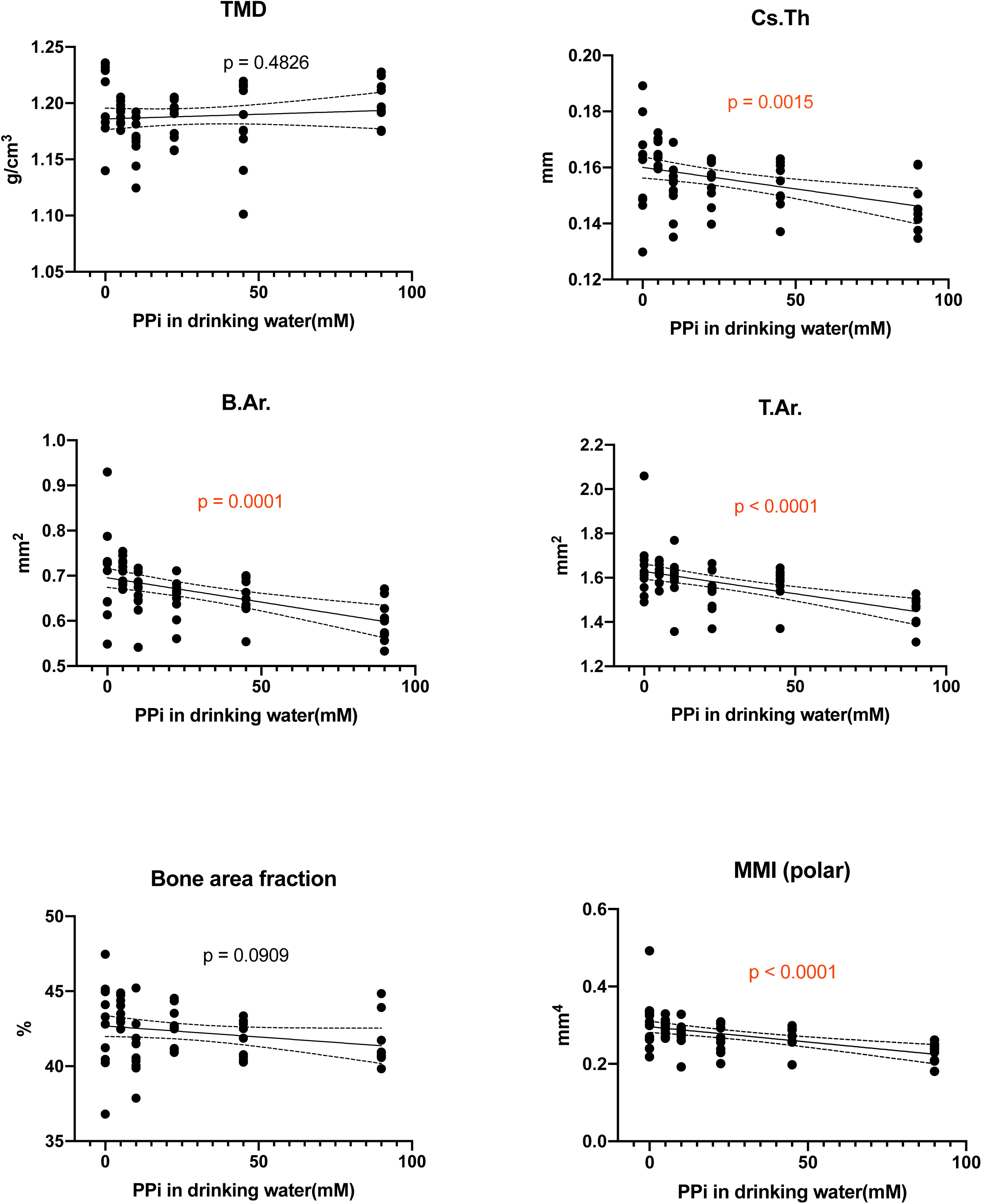
PPi treatment reduced cortical bone quality of femora from Abcc6-/- mice receiving acceleration diet. Various structural parameters of cortical bone were quantified with microCT. TMD (tissue mineral density), Cs.Th (cross sectional thickness), B.Ar. (Bone area), T.Ar. (Tissue area), Bone area fraction (B.Ar./T.Ar.*100), MMI (mean polar moment of inertia). Group trends were quantified with standard regression analysis. Lines represent 95% confidence intervals. Significance was set at p < 0.05, n=10 per group. Dots are individual data points.

## DISCUSSION

In the early 1960’s, Fleisch and colleagues first discovered the ability of PPi to inhibit the complexing of calcium with phosphate molecules to form hydroxyapatite, thereby preventing calcification in soft tissues^9,31^. More recently, patients with PXE were found to have reduced plasma PPi levels which is thought to directly contribute to the progression of ectopic calcification in these patients^15^. Presumably, when PPi is administered to Abcc6-/- mice via the drinking water, PPi levels in the blood plasma are transiently elevated, and calcification in connective tissues that are most susceptible to dysfunction and degeneration is inhibited^16^. Although it is now well established that PPi administration can block ectopic calcification in PXE, not much information is available about the potential side-effects of the high doses of PPi that are needed when administered orally to counteract the ectopic mineralization. In the current study, using an established animal model for PXE, the Abcc6-/- mouse, we 1) addressed the hypothesis that high doses of PPi stimulate ectopic calcification because of the substantially increased intake of Pi. 2) determined the effects of oral PPi administration on bone structural and mechanical properties, and 3) compared the potential of PPi to inhibit aberrant mineral deposition in Abcc6-/- animals receiving standard chow versus a diet high in phosphate and low in magnesium , which is known to stimulate ectopic calcification^23^.

Contrary to our hypothesis, drinking water containing up to 90 mM PPi did not increase the volume of aberrant hydroxyapatite deposition in any of the organs of Abcc6-/- mice analyzed. This finding was also confirmed in Abcc6-/- mice maintained on the acceleration diet. Some loss of structure and strength of the long bones was noted in animals given a high dose of PPi.

In contrast to previous studies much higher doses of PPi were needed to prevent calcification in the eyes and muzzle skin of Abcc6-/- mice. We did not observe a reduction in hydroxyapatite deposition in the eyes or muzzle skin when Abcc6-/- mice were given drinking water with PPi in the range of 5-22 mM, whereas we have previously reported that drinking water with as little as 10 mM PPi almost completely prevented muzzle skin calcification in Abcc6-/- mice^16^. Water may not always be consumed by laboratory animals consistently and at desired volumes^32^, which may partly explain the differences in experimental results seen here, as well as necessitate higher doses of PPi in mice to elicit treatment effects. Secondly, substrains of C57Bl/6J “wild-type” mice with no obvious phenotype but small genetic changes from the parent strain, may be responsible for differences in the uptake and metabolic breakdown of PPi in the intestine^33^. Abcc6-/- mice used in this study were backcrossed for > 10 generations with the parent C57BL/6J strain. Interestingly, studies that have reported inhibition of ectopic calcification in Abcc6-/- mice by providing drinking water containing 10 mM PPi, used animals that were backcrossed for > 15 generations with the C57BL/6J-OLAHSD sub-strain (Theo Gorgels, personal communication)^13,16^. Intriguingly, C57BL/6J- OLAHSD mice harbor biallelic mutations in the genes alpha-synuclein and multimerin-1, resulting in an unexpected reduction of bone mineral density in these animals^33^. Hence, it is conceivable that differences in the regulation of mineral deposition in these C57BL/6J sub-strains underlie at least part of the differences in the effects of orally administered PPi between both Abcc6-/- sub-strains. We plan to test this hypothesis in future experiments. Additionally, differences in rodent diet and the methods used to quantify ectopic calcification, may have contributed to observed differences in the efficacy of low dose PPi treatment. Notably, in this study, we used three dimensional microCT to accurately quantify calcification, whereas Dedinszki et al.^16^ used less quantitative histomorphometry.

Loss of ABCC6 significantly reduced plasma PPi levels, and Abcc6-/- mice showed greatly reduced plasma PPi when compared to their WT counterparts, in line with previous reports^15,34,35^. This result was confirmed in Abcc6-/- mice maintained on standard as well as on acceleration diet. Interestingly, animals on acceleration diet had lower plasma PPi concentrations than animals fed the standard diet. We currently have two potential explanations for these results. First, the amount of PPi in the rodent chow might underlie these differences; standard chow contains ∼85 nmoles PPi/g, whereas the acceleration diet contains ∼5 nmoles PPi/g. The lower amounts of PPi in the acceleration diet may have contributed to the lower plasma PPi concentrations in mice fed this diet. Low levels of PPi in the acceleration diet chow may have contributed to the severe calcification phenotype in Abcc6-/- mice fed this diet. Pomozi et al.^35^ have postulated before that PPi in the food contributes to plasma PPi and subsequent inhibition of ectopic calcification in Abcc6-/- mice. What argues against a major contribution of dietary PPi to plasma concentrations, however, is that we did not detect any increase in plasma PPi in animals receiving drinking water containing 90 mM PPi. Daily intake of PPi via drinking water containing 90 mM PPi is ∼ 1000-fold higher than intake from the standard rodent diet.

An alternative, more likely, hypothesis is that the pro-calcification state in animals fed the acceleration diet, results in more of the plasma PPi being consumed: more PPi in plasma will bind to the increased number of amorphous calcium phosphate nanoprecipitates that we hypothesize will be formed in animals on the acceleration diet. Future studies are needed to elucidate the mechanism underlying the reduced plasma PPi concentrations found in animals receiving the acceleration diet.

ENPP1 is required to generate PPi from ATP, and animals lacking this enzyme exhibit severe ectopic calcification due to the almost complete absence of PPi in the extracellular environment. Mild kidney calcification is evident in 8 weeks old mice lacking ENPP1 activity^16^. Enpp1^asj^ mice that were fed a magnesium enriched diet had markedly reduced mineral deposition in the kidneys at 14 weeks of age^36^. In our studies, we have implemented the reverse strategy of limiting the magnesium content of the diet to stimulate calcification in kidneys that were not visibly affected by loss of Abcc6. As expected, Abcc6-/- mice that were fed the acceleration diet not only showed accelerated muzzle skin calcification, but also developed profound calcification in their kidneys (Fig. 2B). Interestingly, oral PPi completely failed to reduce kidney calcification in Abcc6-/- mice on acceleration diet. Here, the timing of commencement of PPi treatment may also be a factor, since Enpp1^asj^ mice whose mothers consumed 0.3 mM PPi during pregnancy were better responsive to magnesium treatment following birth ^16,36^. It is important to note however, that pre-natal treatments may not be feasible for PXE patients, since this condition is most often diagnosed well after birth, most often in the second decade of life. It is also possible that the dietary magnesium deficiency may have further promoted a pro-calcification environment that is resistant to oral PPi treatment in the kidneys. Further study is required to evaluate the efficacy of PPi treatment for kidney calcification.

PPi has multi-faceted roles in bone. It can bind to mature calcium phosphate crystals and prevent their further growth, but it can also stimulate mitogenic pathways in osteoblasts to promote differentiation and mineralization^37^. Orally administered PPi to some extent accumulates in long bones (Fig. 3). Our finding that femora of Abcc6-/- that were given PPi, demonstrated a reduced ultimate moment (Fig. 4), suggests that PPi accumulation affects bone homeostatic mechanisms, possibly by changing the bone mineral composition. These data are in line with two previous studies reporting loss of trabecular bone structure in the vertebral bone^10^ and in tibiae of aged Abcc6-/- mice (∼2 years)^11^. In the latter study, loss of trabecular bone was not observed in the long bones of 6-month-old Abcc6-/- animals, similar to our results. 3-months-old Abcc6-/- mice maintained on acceleration diet and given oral PPi exhibited more dramatic structural deficits in the femur, particularly decreased cortical bone area and cross-sectional thickness (Fig 7, Supp Fig. 5). Particularly with regard to advanced cases of ectopic calcification typical in aged animals or those subjected to sustained nutritional deficiencies, the effects of oral PPi treatment on the skeleton are subject to further investigation.

Lastly, the authors would like to acknowledge that this study has a few limitations. Firstly, PPi administration via the drinking water may not be feasible for human PXE patients. A more translatable method would be dosing of animals once a day by the oral route. This method is time consuming however, and animals would need to be fed an acceleration diet to limit treatment duration. Secondly, a direct correlation could not be made here between uptake of PPi in the gastrointestinal tract and plasma PPi levels. Hence, changes in plasma PPi as a result of the treatment could not be captured. Thirdly, we did not optimize the formulation of PPi to enhance bioavailability, such as with sodium-free forms of PPi, which may have affected drug pharmacokinetics and by extension its efficacy^38^. Still, oral administration may be convenient for daily PPi administration in PXE patients with their long-term disease prognosis. Possibly, administration in gelatin capsules, as in the ongoing clinical trial in France, enhances palatability and bioavailability^38^.

In conclusion, this study demonstrates that PPi, administered via drinking water over a sustained period of time does not stimulate ectopic calcification in Abcc6-/- mice, and at very high doses prevents calcium phosphate mineral deposition in soft connective tissues like the muzzle skin and eyes. However, the PPi doses effectively blocking disease progression in Abcc6-/- mice here would in humans translate into unrealistic dosage regiments of ∼200g Na_2_PPi/day. While our data do not conclusively demonstrate the suitability of oral PPi administration to prevent disease progression in PXE, they warrant further studies aimed at improving the pharmacodynamics of PPi following its oral administration. This may be achieved by increasing PPi bioavailability or by designing formulations that increase plasma PPi levels to physiological concentrations for multiple hours a day. In addition, it may be prudent to address the variability in the efficacy of orally administered PPi to prevent ectopic calcification in Abcc6-/- mice housed in different laboratories. If a similar variability in the efficacy of oral PPi is present in humans, demonstration of a treatment effect in PXE patients will be challenging. This seems to be an important consideration in the context of the ongoing human clinical trials of PXE. On a positive note, insights to the underlying causes of the substantial variability in the effectiveness of oral PPi administration in translational studies might provide data to improve consistency in treatment outcomes.

## ACKNOWLEDGEMENTS

This research was supported by the National Institutes of Arthritis and Musculoskeletal and Skin Diseases under award nos. R01AR082460 and R21AR083597 to KW. Further funding for this work was provided by PXE International to KW. We thank our colleague Piet Borst (The Netherlands Cancer Institute) for critically reading our manuscript and valuable discussion. The content is solely the responsibility of the authors and does not represent the views of the funding agency.

## AUTHOR CONTRIBUTIONS

IR, NY, JB, CS, SN, DM and FN (Investigation), IR and KW (Writing-original draft), KW (Conceptualization, Funding acquisition), IR, RT and KW (Methodology, Supervision, Formal Analysis, Writing-review and editing).

## DECLARATIONS

The authors declare that they have no known competing financial interests or personal relationships that could have appeared to influence the work reported in this paper.

