## Supplemental Figures for "High-dose oral pyrophosphate inhibits connective tissue calcification in Abcc6 null mice but affects bone structure"

### Supp Fig 1.

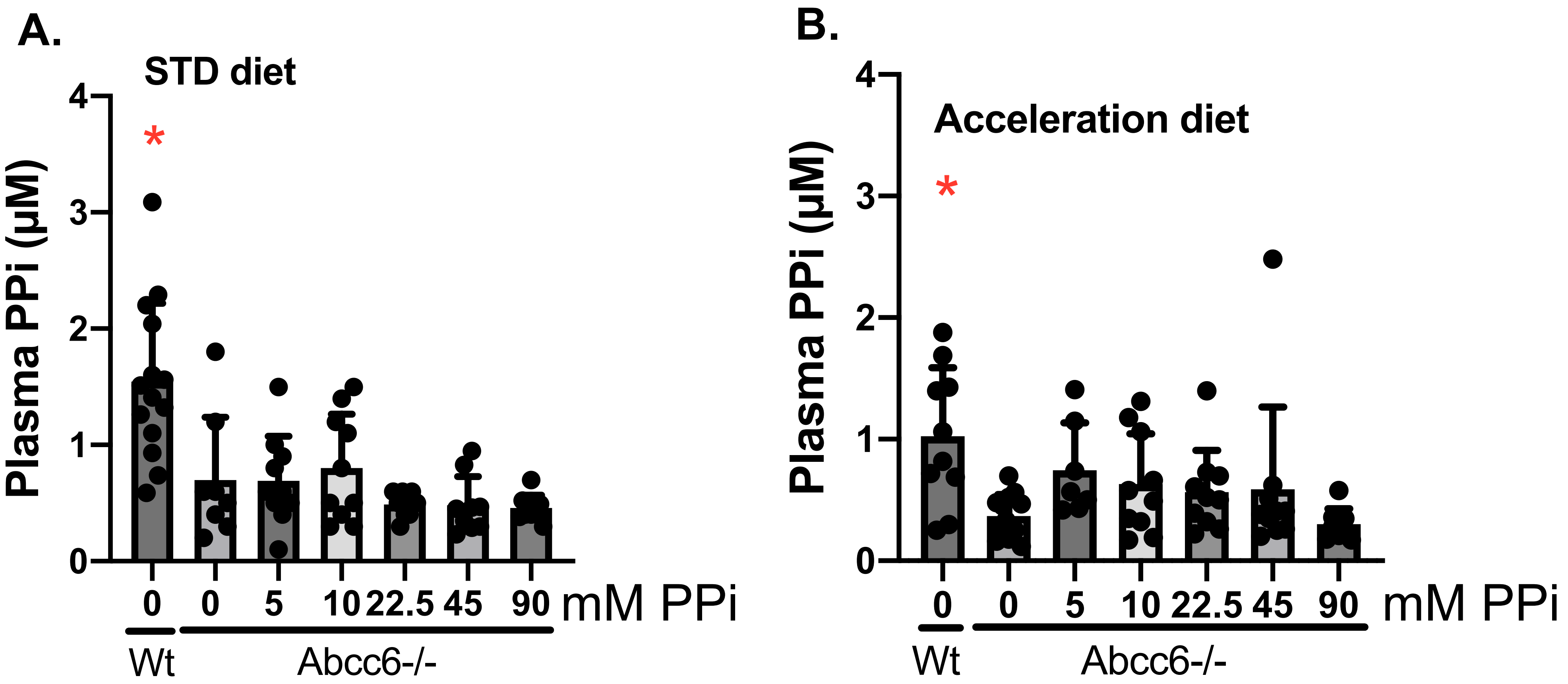
A.
STD diet
Plasma PPi (µM)
0
1
2
3
4
*
0
0
5
10
22.5
45
90
mM PPi
Wt
Abcc6-/-
B.
Acceleration diet
Plasma PPi (µM)
0
1
2
3
4
*
0
0
5
10
22.5
45
90
mM PPi
Wt
Abcc6-/-

**Supp Fig 2. Mechanical and structural properties of femora of wild type and Abcc6-/- mice on standard diet**

**MECHANICAL PROPERTIES**

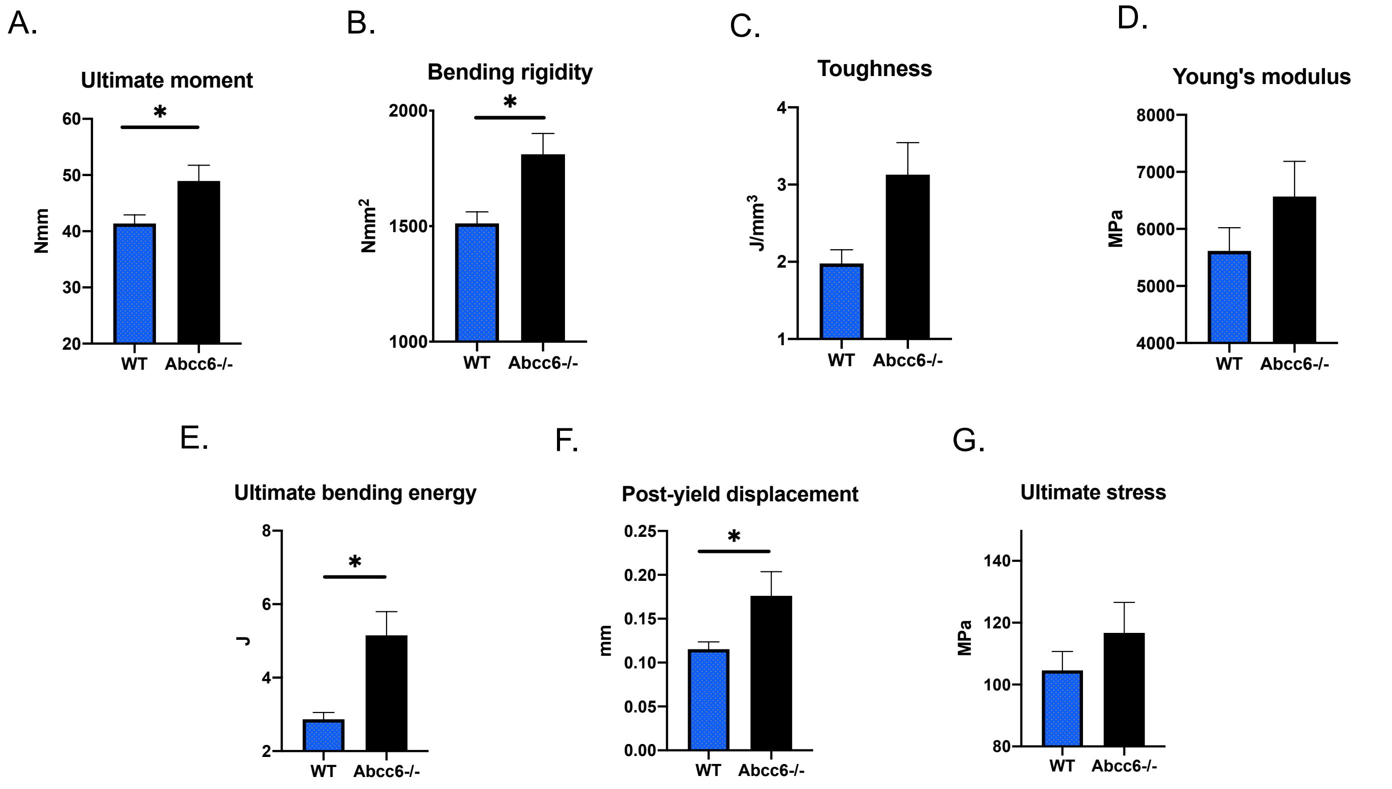


**CORTICAL BONE PROPERTIES**

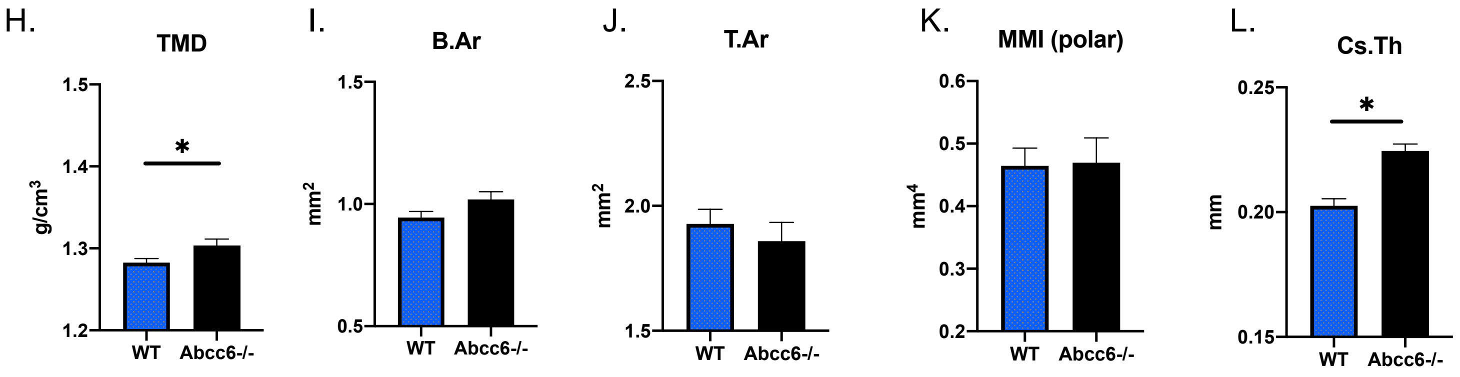


**TRABECULAR BONE PROPERTIES**

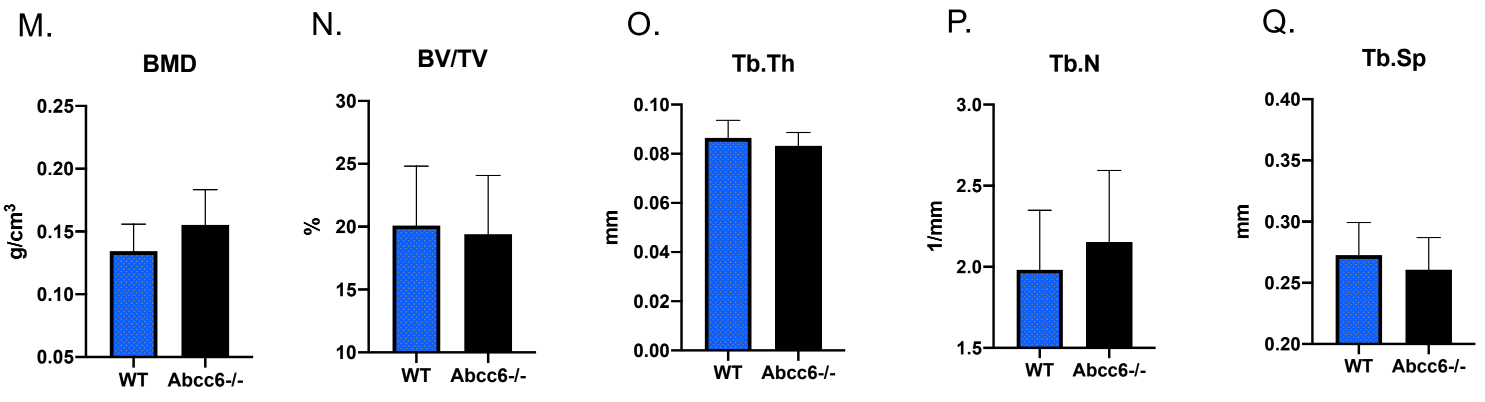

**Supp Fig. 3. Trabecular bone properties of femora of Abcc6-/- mice on standard diet receiving PPi via their drinking water**

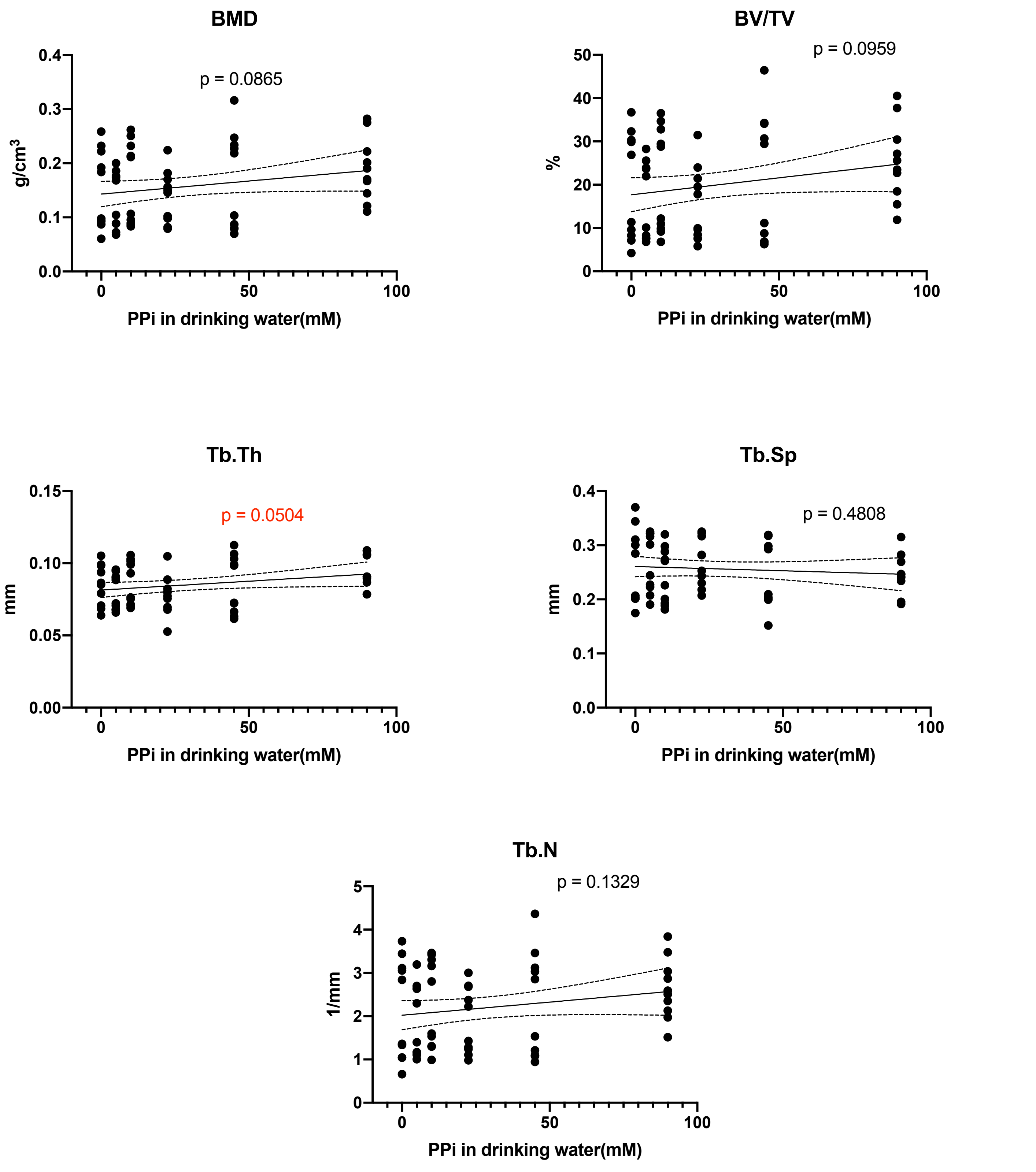

**Supp Fig 4. Mechanical and structural properties of femora of Abcc6-/- mice on acceleration diet**

#### MECHANICAL PROPERTIES

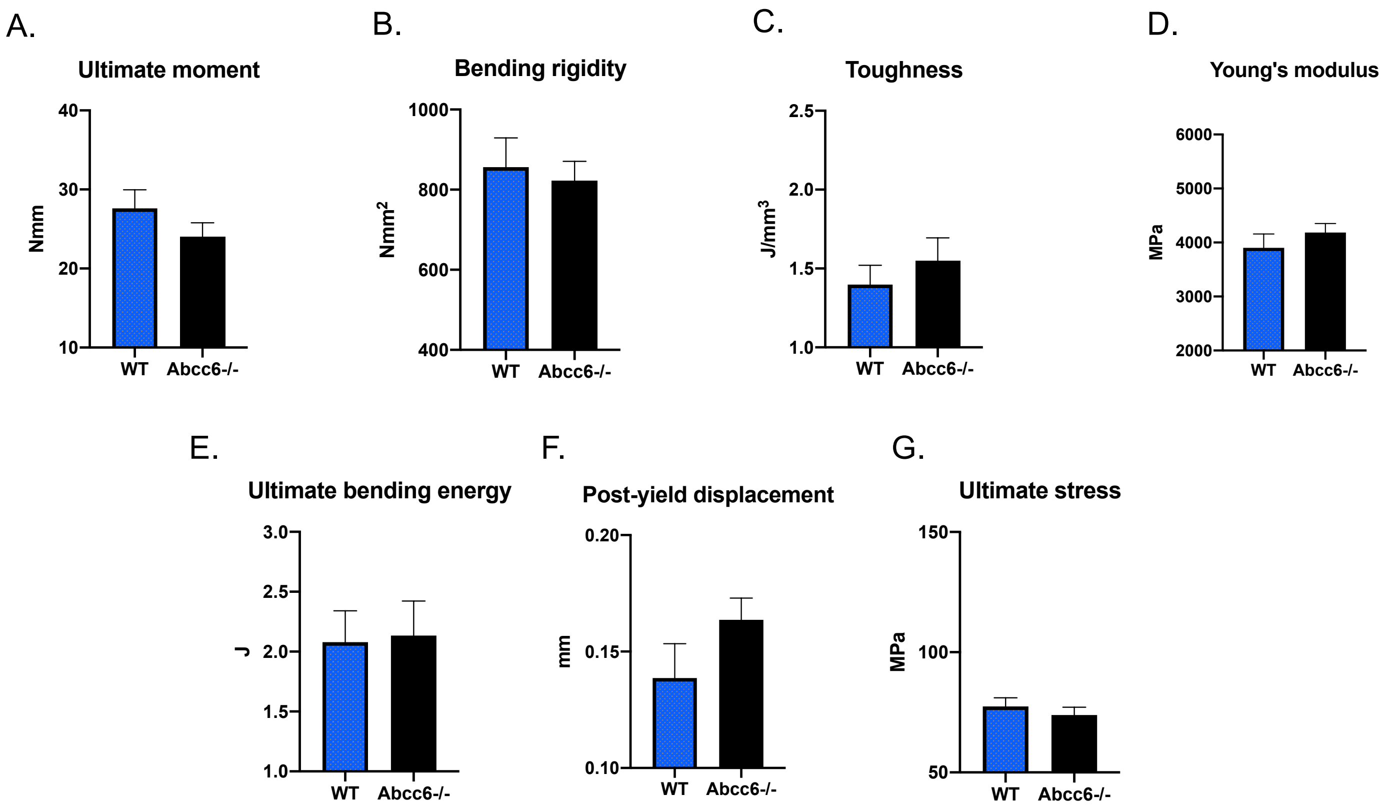


#### CORTICAL BONE PROPERTIES

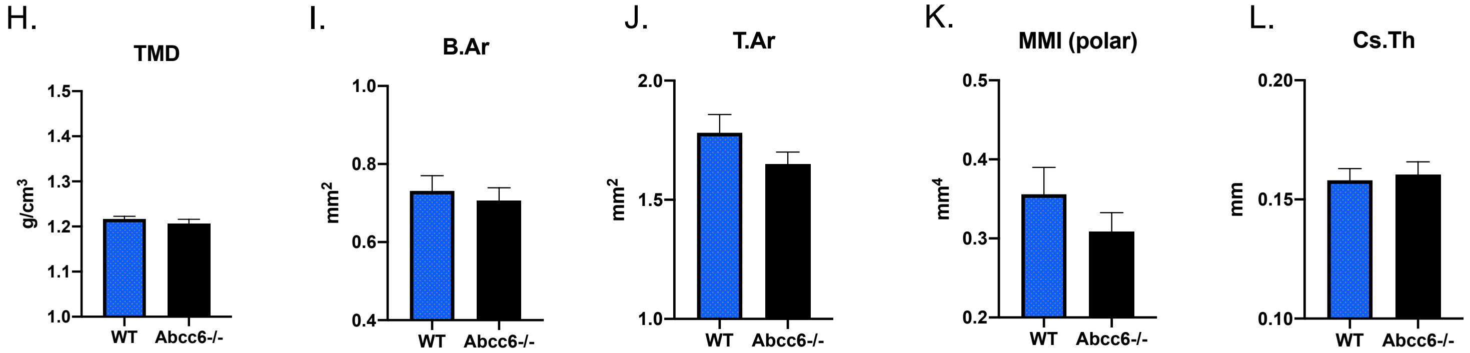


#### TRABECULAR BONE PROPERTIES

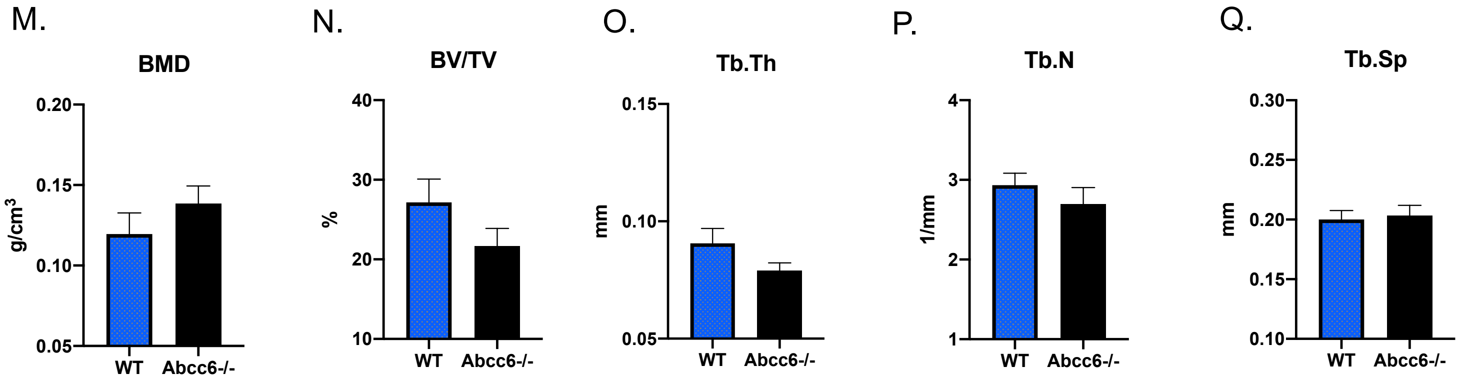

**Supp Fig. 5 Trabecular bone properties of femora of Abcc6-/- mice on acceleration diet receiving PPi via their drinking water**

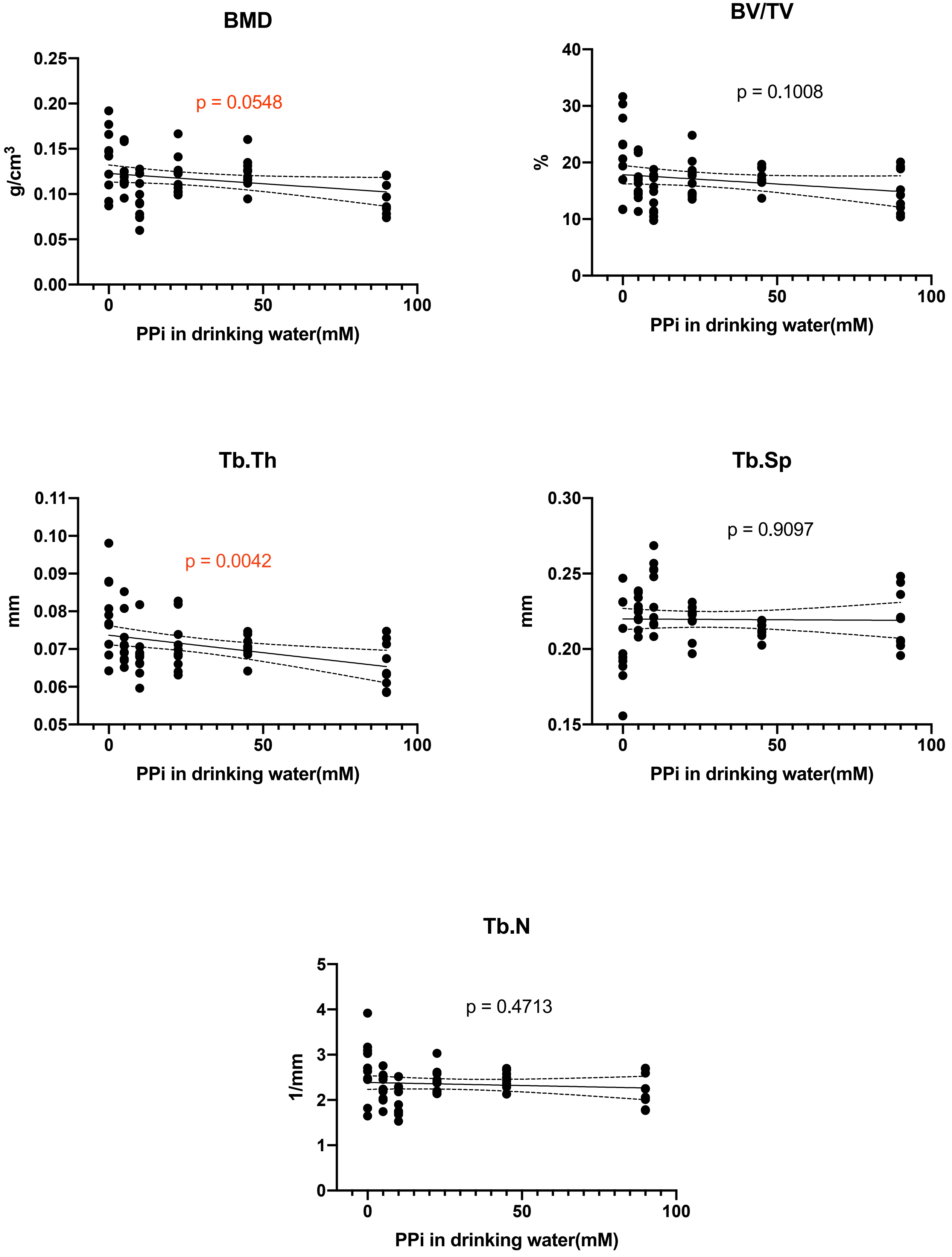

**Legends Supplemental Figures**

**Supplemental Figure 1. Plasma pyrophosphate concentrations in wild type mice and Abcc6-/- mice receiving PPi via their drinking water.** Plasma PPi concentrations were determined in blood samples collected by cardiac puncture at the end of the experiment. Group means were compared using one way ANOVA with post hoc Dunnett's tests. Significance was set at $p < 0.05$, n=10 per group. Error bars represent SD.

**Supplemental Figure 2. Mechanical and structural properties of femora of wild type and Abcc6-/- mice .** Group means were compared with standard t-test. Error bars are mean ± sem. Significance was set at $p < 0.05$, n=10 per group.

**Supplemental Figure 3. Trabecular bone properties of femora of Abcc6-/- mice receiving PPi via their drinking water.** Group trends were quantified with standard regression analysis. BMD (bone mineral density), BV/TV (bone volume), Tb.Th (trabecular thickness), Tb.Sp (trabecular spacing), Tb.N (trabecular number). Lines represent 95% confidence intervals. Significance was set at $p < 0.05$, n=10 per group. Dots are individual data points.

**Supplemental Figure 4. Mechanical and structural properties of femora of wild type and Abcc6-/- mice on acceleration diet receiving PPi via their drinking water.** Group means were compared with standard t-test. Error bars are mean ± sem. Significance was set at $p < 0.05$, n=10 per group.

**Supplemental Figure 5. Trabecular bone properties of femora of Abcc6-/- mice on acceleration diet receiving PPi via their drinking water.** Group trends were quantified with standard regression analysis. BMD (bone mineral density), BV/TV (bone volume), Tb.Th (trabecular thickness), Tb.Sp (trabecular spacing), Tb.N (trabecular number). Lines represent 95% confidence intervals. Significance was set at $p < 0.05$, n=10 per group. Dots are individual data points.
